# *OPEN leaf*: an open-source cloud-based phenotyping system for tracking dynamic changes at leaf-specific resolution in Arabidopsis

**DOI:** 10.1101/2021.12.17.472861

**Authors:** Landon G. Swartz, Suxing Liu, Drew Dahlquist, Skyler Kramer, Emily S. Walter, Sam McInturf, Alexander Bucksch, David G. Mendoza-Cozatl

**Affiliations:** Department of Electrical Engineering and Computer Science, University of Missouri-Columbia, USA; Department of Plant Science and Technology, College of Agriculture, Food, and Natural Resources, University of Missouri, Columbia, USA; Department of Plant Biology, University of Georgia, Athens, GA 30602, Warnell School of Forestry and Natural Resources, University of Georgia, Athens, and Institute of Bioinformatics, University of Georgia, Athens, GA 30602; MU Institute of Data Science and Informatics, University of Missouri-Columbia, USA

**Keywords:** CyVerse, High-throughput phenotyping, hydroponics, phenomics, Plant nutrition, *Arabidopsis thaliana*, Internet of Phenotyping

## Abstract

The first draft of the *Arabidopsis* genome was released more than 20 years ago and despite intensive molecular research, more than 30% of *Arabidopsis* genes remained uncharacterized or without an assigned function. This is in part due to gene redundancy within gene families or the essential nature of genes, where their deletion results in lethality (*i.e*., the *dark genome*).

High-throughput plant phenotyping (HTPP) offers an automated and unbiased approach to characterize subtle or transient phenotypes resulting from gene redundancy or inducible gene silencing; however, commercial HTPP platforms remain unaffordable. Here we describe the design and implementation of *OPEN leaf*, an open-source HTPP system with cloud connectivity and remote bilateral communication to facilitate data collection, sharing and processing.

*OPEN leaf*, coupled with the SMART imaging processing package was able to consistently document and quantify dynamic morphological changes over time at the whole rosette level and also at leaf-specific resolution when plants experienced changes in nutrient availability.

The modular design of *OPEN leaf* allows for additional sensor integration. Notably, our data demonstrate that VIS sensors remain underutilized and can be used in high-throughput screens to identify characterize previously unidentified phenotypes in a leaf-specific manner.

**Significance Statement:** Many bottlenecks exist in high-throughput phenotyping involving computing power for processing and a lack of focus on abiotic stresses that has prevented an advancement in phenotyping on par with genotyping. Therefore, we create an automated HTP system that performs nutrient studies on *Arabidopsis thaliana* with cloud-based image processing that quantifies plant traits at a whole and leaf-level.

## Introduction

Understanding genome-to-phenome relationships is at the core of *predictive biology*, whose aim is to predict a biological outcome from a known input (Lopatkin & Collins, 2020). The phenome, however, is also the result of specific environmental conditions and the ability of organisms to sense and adapt to changes in their environment (Des Marais *et al.*, 2013; Xu, 2016; Buckner *et al.*, 2019). Thus, *predictive biology* requires deep mechanistic understanding of the genetic makeup of a given organism but also its plastic response to environmental cues. In the case of plants, these cues may include changes in light intensity, nutrient, and water availability, and even the presence of other organisms, including microbes or additional plant species (Lundberg *et al.*, 2012; Legris *et al.*, 2016). While significant advances have been made to understand plant behavior at the molecular, physiological, and ecological level, the prediction of plant responses during changing environments remains a formidable challenge that requires consistent collection of reproducible data at the “*-omic*” and environmental levels.

Collecting large amounts of genomic and environmental data have become reasonably affordable in recent years; however, integrating, sharing, and analyzing such data in real-time is currently a bottleneck in biology. Moreover, the methods for characterizing genomes and phenomes have not advanced at the same rate (Moore *et al.*, 2013; Yang *et al.*, 2020). While genomic techniques have benefited from new developments in DNA sequencing, including higher resolution and lower costs, plant phenotyping has improved at a slower rate and commercial plant phenotyping platforms are still costly and practically inaccessible for the majority of plant laboratories (Shendure & Ji, 2008; Jackson *et al.*, 2011; White *et al.*, 2012; Reynolds *et al.*, 2019). More importantly, this gap between genomic and phenomic technologies are preventing the full use of resources available in several plant species such as mutant collections or diversity panels to pursue genome-wide studies or the characterization of complex traits at a higher resolution (Xiao *et al.*, 2017). At the same time, the technology needed for high-throughput plant phenotyping studies including sensors, computer vision, and information technology (IT), have become more accessible, allowing the development of diverse platforms for plant phenotyping experiments including roots (Le Marié *et al.*, 2014; Mathieu *et al.*, 2015), shoots (Awlia *et al.*, 2016; Flood *et al.*, 2016), plant growth within laboratory or greenhouse spaces (van der Heijden *et al.*, 2012; Yang *et al.*, 2014) and large field crops (Sadeghi-Tehran *et al.*, 2017; Maes & Steppe, 2019). Aside from biological data, these platforms have also identified three major limitations in high-throughput plant phenotyping (HTPP) platforms, that is: (i) the ability to consistently reproduce abiotic stresses such as water and nutrient limitation (ii) data management that is standardized and comprehensive for collaborative analysis and re-analysis (iii) and the high cost of commercial high-throughput phenotyping machines (Billiau *et al.*, 2012; Tsaftaris & Scharr, 2019; Reynolds *et al.*, 2019; Yang *et al.*, 2020).

Here we describe *OPEN leaf* [Open PhENotyper for *leaf* **tracking**], an open-source plant phenotyping platform designed to track growth and development of rosette leaves in a leaf-specific manner. Our system was built considering accessible materials worldwide and it is primarily based on a high-resolution camera hovering over user-defined positions to track the morphology (size, shape, color) of the whole rosette and specific leaves. As a semi-autonomous platform, *OPEN leaf* is enabled to communicate remotely with users on a user-defined basis to track data in real-time, thus allowing adjustments throughout the experiments. *OPEN leaf* was developed to be modular, scalable, and includes cloud-based capabilities for data sharing and processing. As a proof-of-concept, here we describe the use of *OPEN leaf* to characterize *Arabidopsis* plants grown hydroponically with different nutrient levels. Reproducibility in the field of plant nutrition is a major roadblock, particularly in micronutrient deficiency studies, where the absolute concentration of elements in the nutrient solution and their ratio with other micronutrients have profound effects on day-to-day phenotypes (Nguyen *et al.*, 2016). With its cloud-based capabilities enabled, *OPEN leaf* was able to resolve dynamic phenotypes in a leaf-specific resolution during changes iron/zinc ratios. Our results demonstrate that within the *Arabidopsis* rosette, multiple phenotypes remain to be identified and that clear developmental changes occur in parallel to the standard chlorosis induced by iron deficiency.

## Materials and Methods

### Hardware

*OPEN leaf* was built using the open-source T-slot aluminum beams from 80/20 (https://8020.net) and a C-Beam Linear Actuator Bundle from OpenBuilds, which contains the track, a gantry plate, and a NEMA23 stepper motor (https://openbuildspartstore.com/c-beam-linear-actuator-bundle/). A high-resolution camera (Allied Vision Mako G-503) was mounted on the gantry plate using a bracket 3D printed on a Prusa i3 MK3 (STL file provided in Supp data; https://www.prusa3d.com). The track system moves to pre-determined positions using GRBL, an open-source, embedded, and high-performance g-code parser controlled by an Arduino microcontroller (https://www.arduino.cc). For a detailed bill of materials, 3D files, schematics and assembly manual see *Supplementary data*.

### Software

OPEN *leaf* is controlled by an open-source desktop application written in C# (*OPEN Controller*, github.com/DMendozaLab/OPEN-Controller), and allows multi-machine control. From the desktop application, the user can identify and select the proper controller (machine) and its unique camera, or other sensors associated with it. The user also selects pre-defined camera settings including gain and aperture, the folder where the pictures will be saved and the GRBL commands to move the gantry plate with the camera attached to specified locations on the track. At each specific location, the camera will take a picture at one time point or a time series for a period of time defined by the user. Installation of *OPEN Controller* is streamlined by a package (*i.e*., installer), which downloads and compiles all the required dependencies.

### Cloud Integration

Integration for CyVerse (https://CyVerse.org) is an optional module of the OPEN series of devices that will send data to CyVerse for cloud storage, access, management, sharing, and analysis. CyVerse integration is done through the desktop application Cyberduck and a command-line interface (CLI) that interacts with CyVerse. Using CyVerse configuration profiles, one can log into a desired CyVerse account to have remote access to a respective CyVerse Data Store. For automated syncing (uploading), we created a python script titled *cyverse.py* that runs a duck - synchronize command to synchronize all data in the local directory to the remote directory located on CyVerse.

### Slack Integration

We used the Slack tool called “bot” account to relay messages regarding the operational status of our OPEN machines. The “bot” account is setup onto a desired channel within a Slack teamspace with the proper permissions. Instructions to integrate Slack with OPEN machines can be found on (github.com/DMendozaLab/OPEN-leaf-cloud). For messages, we first query whether the HTP machine is currently running or not. If the machine is running then a message of “Status: Running” is sent with the current experiment name, and the latest single cycle images are sent. If the machine is not running, then a message of “Status: Dormant” is sent. After seven dormant messages are sent, no more dormant messages are sent until a new experiment is started and the status is updated to “Running”. The status of the machine can be scheduled at specific times per day or “on demand” using slash commands (*e.g*., /srp-get-status).

### Image processing using *SMART*

*SMART* (Speedy Measurement of Arabidopsis Rosette Traits) is a complete set of parameter-free pipeline to analyze plant images in the visible (VIS) and near-infrared spectrum (NIR). The SMART pipeline use top-view images of *Arabidopsis* to compute geometrical traits of the whole rosette and if desired, SMART can also extract information from individual leaves in within the rosette (Table 1). For plant object segmentation and unlike other plant image pipelines that use a threshold value in color space, SMART uses a dominant color clustering method to segment the plant object from the background. The pipeline used are available as source code on GitHub (https://github.com/Computational-Plant-Science/SMART) and are also available as a prepackaged Docker container (https://hub.docker.com/r/computationalplantscience/smart). For the identification and characterization of individual leaves and their corresponding parameters, SMART adopted a watershed segmentation algorithm. This algorithm first uses Euclidean Distance Transform to compute the Euclidean distance map to the closest zero (*i.e*., background pixel) for each of the foreground pixels. Then each peak in the Euclidean distance map was fed as the markers for watershed segmentation to segment and count each of the plant leaves (van der Walt *et al.*, 2014). In addition, SMART was able to compute the color distribution and recognize distinct shades of color on the plant leaf surface.

**Table 1.** Summary of plant traits measurement and units

| Measurement | Units | Trait type* | Description |
| --- | --- | --- | --- |
| Whole plant area | pixels | WP | Sum of the whole plant area |
| Whole plant width/height | pixels | WP | The width/height of the plant object, computed from major/minor axis values of the fitted bounding ellipse. |
| Whole plant solidity | None | WP | The ratio of fitted whole plant contour area to its convex hull area, describe the shape of the whole plant, A value of 1 signifies a solid object and a value less than 1 will signify an object having an irregular boundary. |
| Whole plant curvature | None | WP | The derivative of the inclination of the tangent with respect to arc length. The mean curvature value of all leaves, describing the boundary shape change of individual leaf. |
| Number of leaves | None | WP | Count of the total number of leaves |
| Color distribution | None | LS | The distribution and percentage of distinct shades of color on the plant leaf surface. |
| Leaf area | pixels | LS | Individual leaf area |
| Leaf width/height | pixels | LS | The width/height of the individual leaf object, computed from major/minor axis values of the fitted bounding ellipse. |
| Leaf solidity | None | LS | The ratio of fitted leaf contour area to its convex hull area. |

\* WP, whole plant; LS, Leaf specific.

### Plant growth conditions

Arabidopsis Col-0 seeds were surfaced sterilized and geminated on 1/4 Murashige and Skoog (MS) medium after stratification for 2-days at 4°C in the dark. Then, seeds were allowed to germinate under 16 h light/ 8h dark cycles, 23°C day/ 21°C night (60 % humidity) as previously described (Nguyen *et al.*, 2016). After the emerging of the first true leaves (10-12 days after seed plating), plants were transferred to black magenta boxes using 3D printed floaters (See Supp data for STL files). Unless otherwise stated, plants were maintained in standard Arabidopsis hydroponic media as described in Nguyen et al. (2016) and imaged every four hours using the *OPEN leaf* system described here. Hydroponic media were saturated with air using standard aquarium air pumps and bubbling stones and the nutrient solution was changed every three days. After 4-6 days of continuous image capturing, the nutrient solution of some plants was replaced with hydroponic without Fe or with hydroponic media containing 10 μM Zn with and without Fe (the standard Zn concentration was 1 μM Zn).

## Results

### OPEN leaf design

High-throughput plant phenotyping (HTPP) has revolutionized the way plant scientists gather, process and extract information from plants, including model and crop species.. However, commercial HTPP stations remain exceptionally expensive for laboratories around the world. Alternatively, several open-source HTPP devices are now available, but their assembly and operation still require a high degree of proficiency in mechanical engineering or computer sciences. To address this issue, we designed a top-view single row phenotyper (*OPEN leaf*; Figure 1A). We streamlined the assembly of *OPEN leaf* by taking advantage of affordable pre-assembled tracks available worldwide combined with controllers and open-source software (Figure 1B). At the core of *OPEN leaf* is a high-resolution camera that captures images suitable to be processed with current computer vision pipelines. In addition, we took advantage of CyVerse, a publicly accessible cyberinfrastructure, and Slack, a team communication platform, to enable cloud-storage, data sharing and processing, and a two-way communication protocol to remotely request operational status and data from *OPEN leaf* on a regularly basis or on-demand (Figure 1C).

**Figure 1.**
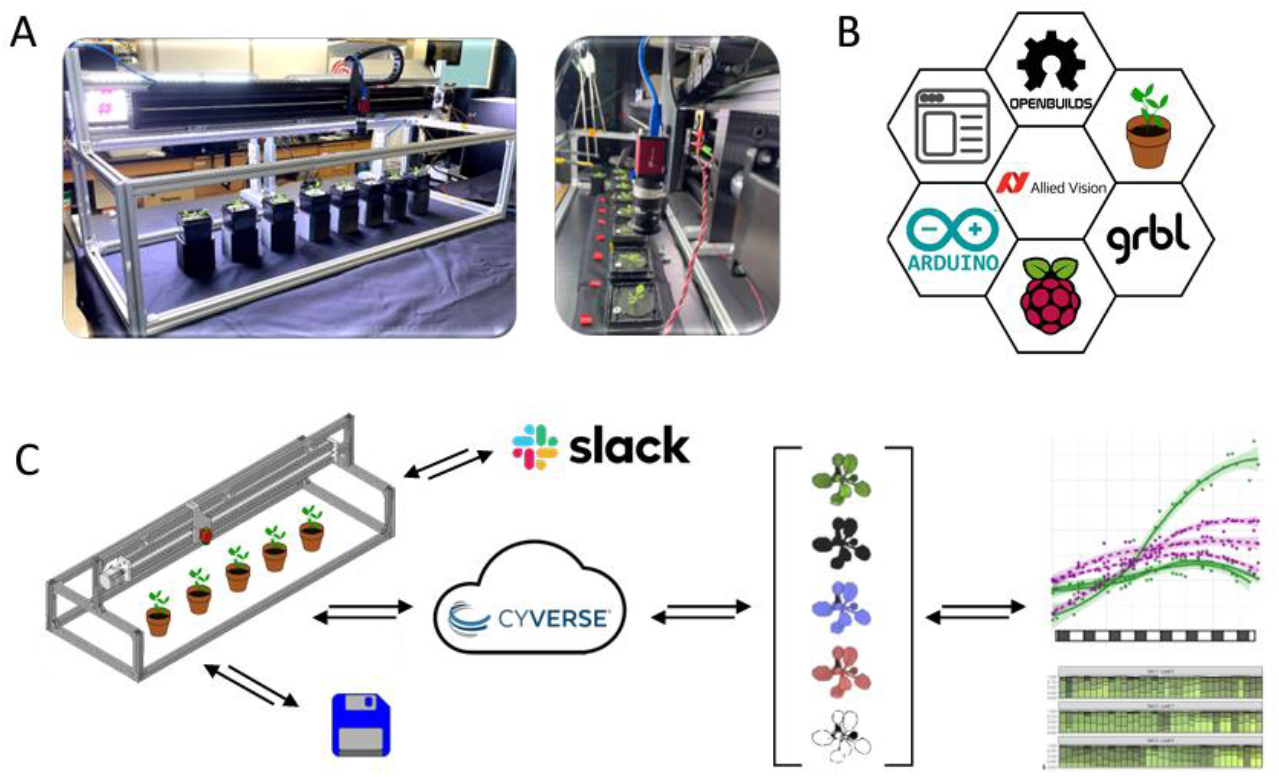
*OPEN leaf* design and capabilities. (a) The main components of the OPEN *leaf* imaging system are aluminum extrusion beans, a linear actuator and a high-resolution VIS camera to capture top-view images of Arabidopsis plants grown hydroponically. (b) The software and hardware required to operate OPEN *leaf* is based on publicly available packages and materials available worldwide. (c) To facilitate data management, we have included optional cloud-based connectivity for remote data sharing processing and real-time operational status updates initiated on-demand by the end user.

### OPEN leaf user interface and remote connectivity

The *OPEN leaf* user interface (*OPEN controller*) was designed so users can easily identify the critical parts of the system (*i.e*., camera, storage path, and coordinates to stop along the track; Figure 2A) and have a one-stop access to modify each of them as needed. *OPEN controller* also allows users to capture images at one time point (single cycle) or over a user-defined period of time (time lapse). Images are initially stored locally in a hard drive, but we have also taken advantage of the publicly available cloud cyberinfrastructure CyVerse and its *Discovery Environment* to upload images regularly and facilitate data sharing and processing (scripts available in Supplementary Data). To further engage remote access and communication with *OPEN leaf*, we implemented a two-way communication using Slack (Figure 2B, scripts also available in Supplementary Data). This remote communication allows regularly updates on the operational status of the machine together with the last set of images taken when an experiment is ongoing. Alternatively, if the machine has finished the required cycles or the machine has stopped working due to a power interruption or computer malfunction the user will receive “dormant” status at the specified time (Figure 2B-C). Operational status can also be requested on demand with a similar outcome: “running + experiment + data” or “dormant” (Figure 2D-E).

**Figure 2.**
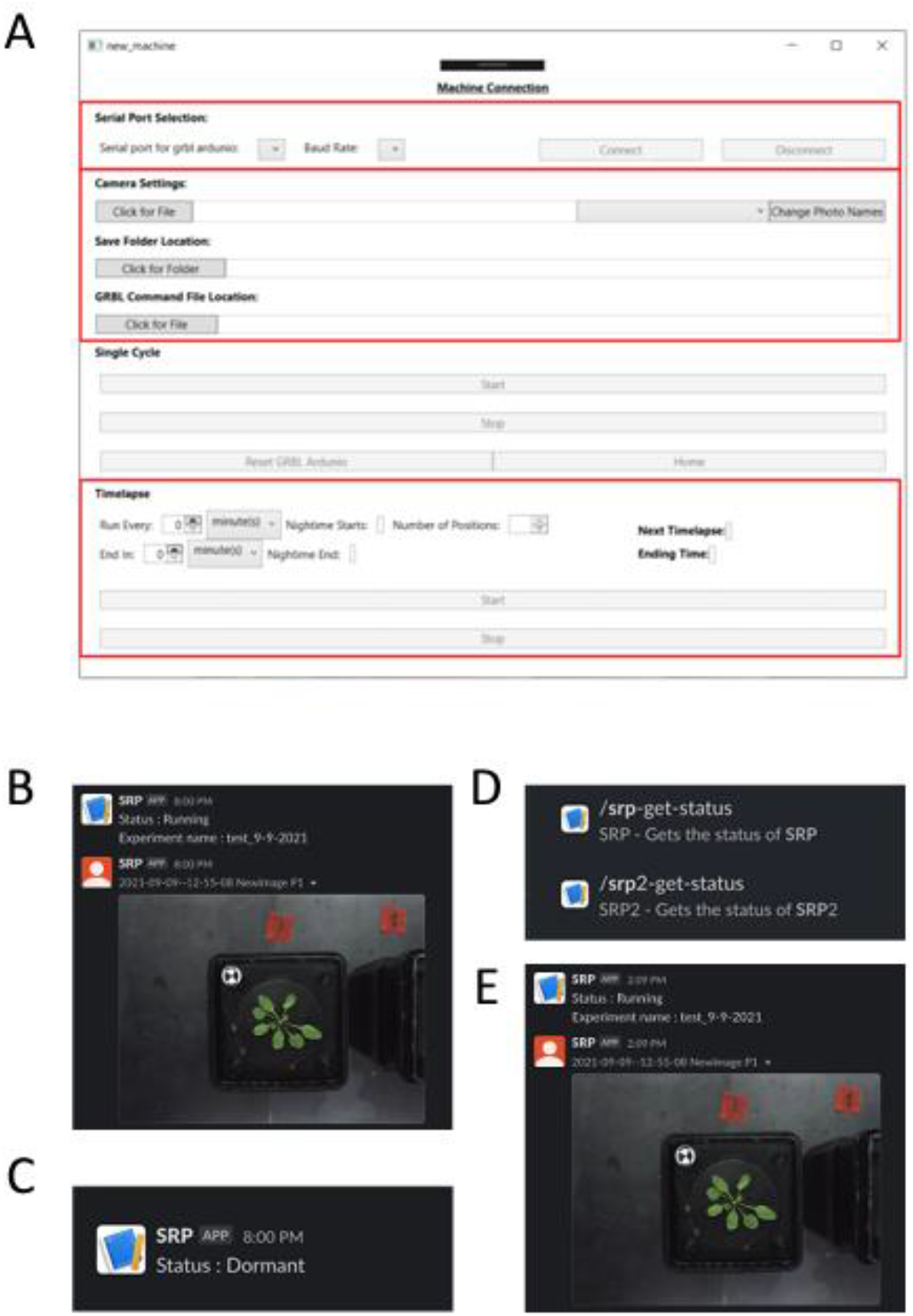
*OPEN* leaf user interface and remote communication. (a) The OPEN *leaf* user-interface was designed to be linear and intuitive with one-to-one relationships to control the different settings of the imaging system. (b-c) By enabling the communication platform Slack, users can receive automatic updates of ongoing experiments indicating the status of the machine and the latest acquired data. (c) In addition, users can request on-demand updates for independent machines running simultaneously.

### Feature extraction of rosette leaves using SMART

To begin testing the capabilities of *OPEN leaf* and the SMART image processing package, we first tracked the growth of *Arabidopsis* rosettes grown hydroponically in replete media or lacking Fe for 7 days. In these experiments, plants were first germinated on agar plates for 7-10 days before being transferred to magenta boxes with replete nutrient solution. After two days of acclimation in the hydroponic system, plants were placed under the *OPEN leaf* track and imaged continuously for two weeks. During the first week plants were grown in replete media to capture the baseline growth and after a week, half of the plants were grown on hydroponic media lacking Fe. While the experiment was running, images were uploaded to CyVerse and processed using the *Speedy Measurement of Arabidopsis Rosette Traits* (SMART) package (Figure 3A). SMART uses the top view of the *Arabidopsis* rosette to extract several geometrical traits of the whole rosette (Table 1). Figure 3B shows an example of the automated rosette segmentation of plants grown in replete media or Fe deficiency. Only 9 representative images throughout the entire experiment are shown but all traits were calculated at each experimental time point. Table 2 shows additional traits calculated for these 9 samples and the full table containing all the data points and traits, including color clustering, is available as Supplementary Table 1. Figure 3C shows the area of the entire rosette of plants grown in the presence and absence of Fe and illustrates how Fe deficiency has a severe negative impact on growth and development, here measured as full rosette area. Note that the sensor used in these experiments was a visible image sensor (*i.e*., a RGB color camera), As such, data collection was only possible during the light cycle of the imposed photoperiod (shown as open blocks on the *x* axis of Figure 3C). The selection of a visible image sensor was selected to maintain an affordable cost with the highest resolution possible and allowed us to uncover novel phenotypes (see below); however, other sensors such as NIR cameras are compatible with the *OPEN leaf* track system and the SMART imaging software.

**Table 2.** Main traits extracted from Arabidopsis plants grown hydroponically shown in Figure 3B.

| Time<br>(min) | Area |  | Solidity |  | Max_width |  | Max_height |  | Curvature |  | Leaf number |  |
| --- | --- | --- | --- | --- | --- | --- | --- | --- | --- | --- | --- | --- |
|  | Full | -Fe | Full | -Fe | Full | -Fe | Full | -Fe | Full | -Fe | Full | -Fe |
| 72 | 29.41 | 33.10 | 0.571 | 0.611 | 211 | 288 | 302 | 269 | 0.026 | 0.022 | 6 | 5 |
| 102 | 43.19 | 51.08 | 0.547 | 0.588 | 285 | 371 | 370 | 324 | 0.022 | 0.020 | 7 | 6 |
| 126 | 58.25 | 72.87 | 0.520 | 0.552 | 360 | 463 | 416 | 426 | 0.017 | 0.018 | 7 | 7 |
| 168 | 104.2 | 109.1 | 0.478 | 0.488 | 626 | 539 | 544 | 636 | 0.017 | 0.019 | 9 | 10 |
| 189 | 123.6 | 129.1 | 0.484 | 0.512 | 718 | 538 | 576 | 658 | 0.016 | 0.017 | 10 | 11 |
| 213 | 143.7 | 156.0 | 0.482 | 0.493 | 808 | 666 | 596 | 731 | 0.017 | 0.016 | 11 | 12 |
| 237 | 161.9 | 170.2 | 0.542 | 0.432 | 651 | 818 | 620 | 792 | 0.012 | 0.014 | 8 | 11 |
| 261 | 237.9 | 200.8 | 0.509 | 0.436 | 963 | 891 | 782 | 816 | 0.013 | 0.014 | 11 | 12 |
| 295 | 290.8 | 257.3 | 0.454 | 0.431 | 1089 | 982 | 1001 | 936 | 0.014 | 0.013 | 11 | 10 |

\*See Table 1 for units.

**Figure 3.**
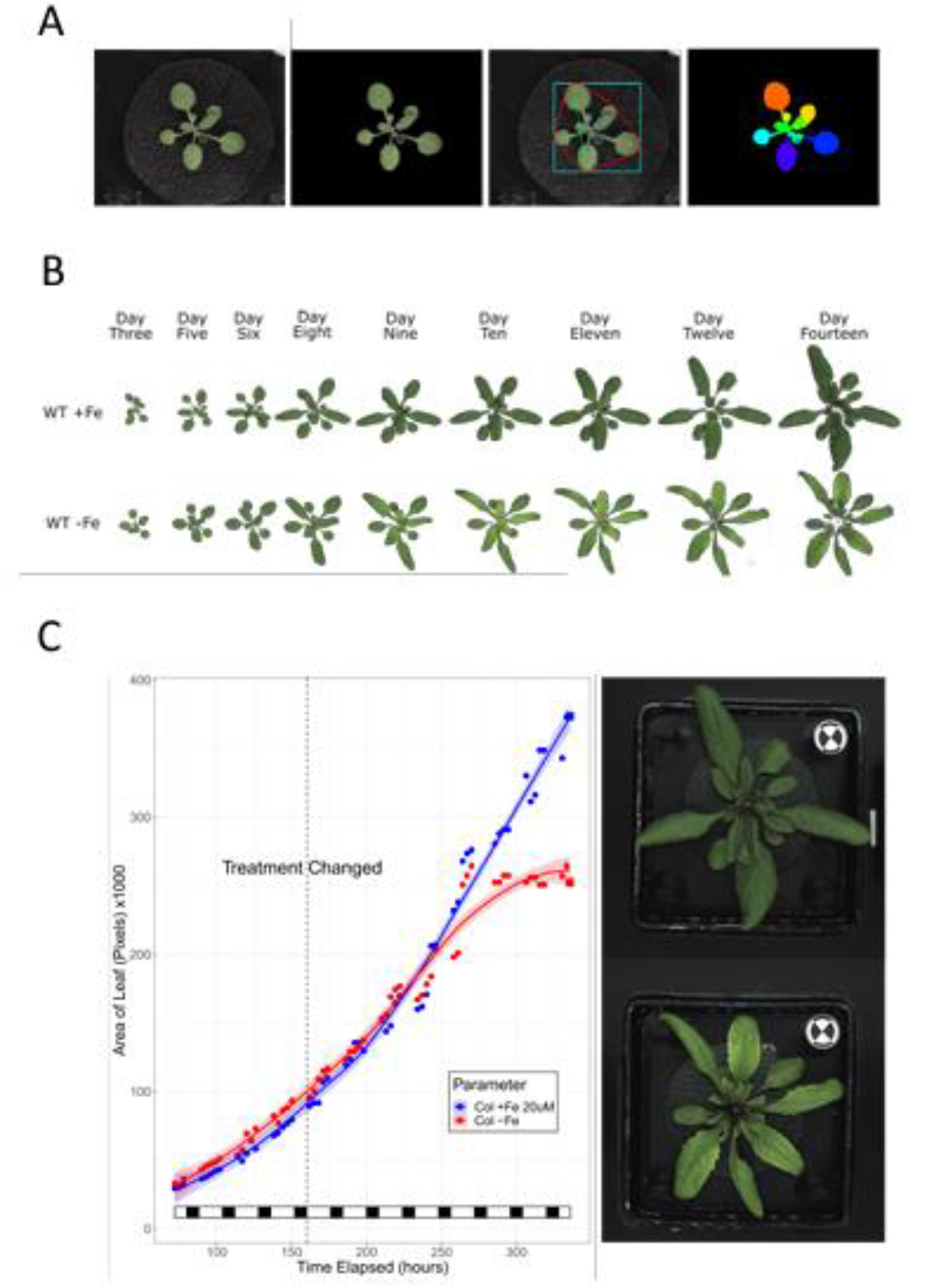
SMART imaging capabilities. (a) The *Speedy Measurement of Arabidopsis Rosette Traits* (SMART) is an automated pipeline to extract morphological information from Arabidopsis plants grown on the OPEN *leaf* imaging system, (b) the SMART pipeline can segment the Arabidopsis rosette and quantify several traits including leaf number, area and color [for a full list of traits see Table 1]. (c) OPEN *leaf* and SMART are capable of detecting visual changes of Arabidopsis plants grown in the presence or absence of iron in the growth media. The figure shows a representative experiment and similar trends were observed in three independent experiments.

Next, we tested whether our phenotyping platform was sensitive enough to detect changes when plants experience a combination of micronutrient availability. In these experiments, plants were grown hydroponically as described before but the nutrient challenge consisted in Fe deficiency experiments in the presence of standard zinc (Zn) concentrations (1 μM) or a mild elevated yet non-lethal concentration of Zn (10 μM) in the hydroponic solution. Figure 4A shows representative rosette images of each condition extracted using the SMART package and Figure 4B shows the area of the whole rosette plotted against time. Both, the images and the corresponding rosette, were able to capture differences in growth and development in each of the growth conditions. Notably, the addition of 10 μM Zn to the hydroponic media resulted in a smaller rosette area compared to plants grown in 1 μM Zn grown with replete Fe (1 μM Zn+Fe) or exposed to Fe deficiency (1 μM Zn-Fe). Moreover, the color clustering algorithm revealed clear differences in color distribution across treatments, a trait that was found to be independent of rosette size and therefore should be extracted and analyzed in parallel. That is, the rosette area of plants grown in 1 μM Zn-Fe was larger than plants grown in 10 μM Zn+Fe, but the color distribution reported as hexadecimal values was consistent with chlorosis only in plants grown without Fe (Figure 4B).

**Figure 4.**
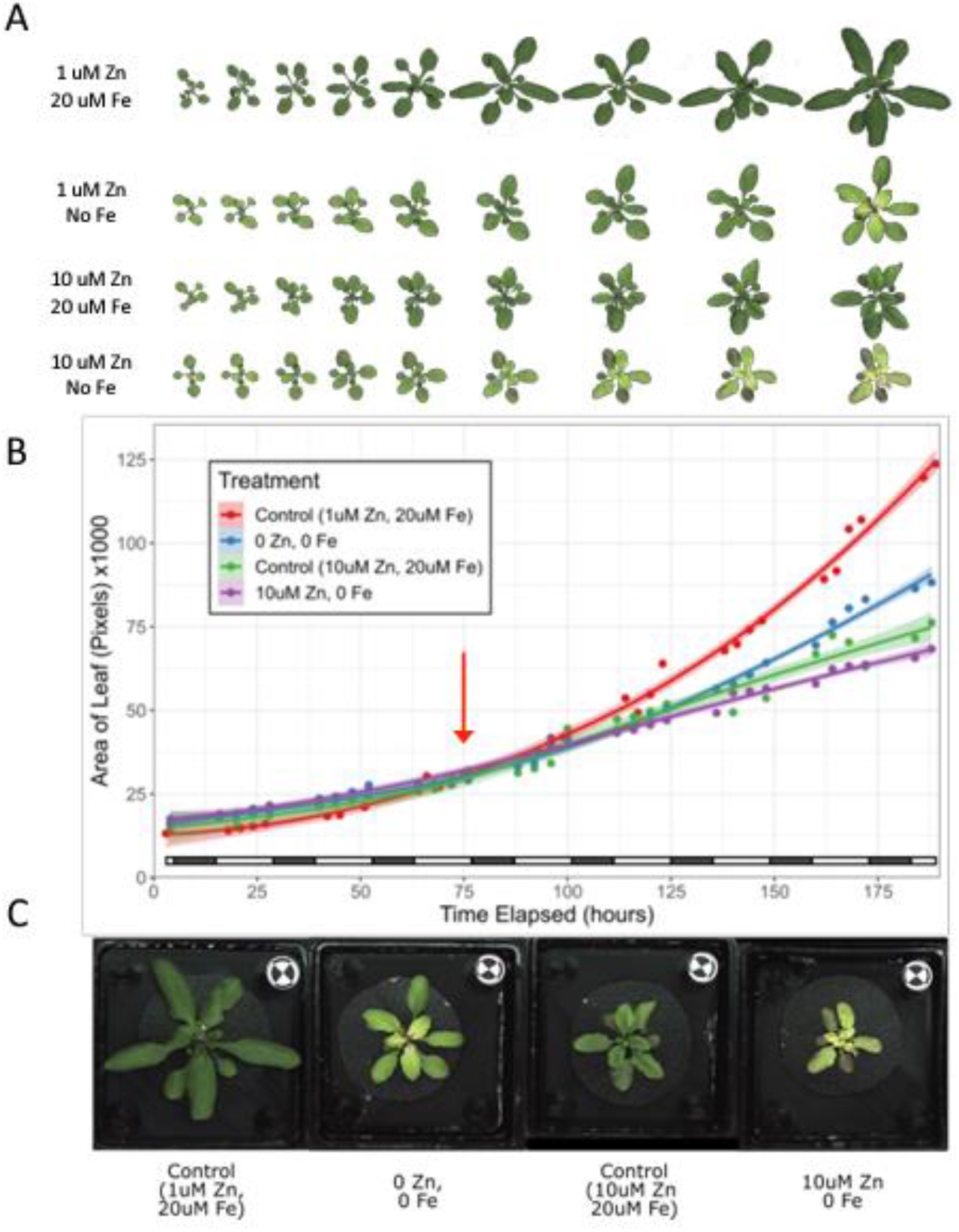
Arabidopsis phenotyping during contrasting changes in nutrient availability. (a) Whole rosette area and color are affected when the ratio of iron (Fe) and zinc (Zn) are modified in the hydroponic media. (b) the *OPEN leaf*-SMART pipeline was able to detect subtle changes in rosette area and color when the Fe/Zn ratio was modified. (c) Changes in Fe and Zn lead to unique phenotypes in individual leaves within the entire rosette in a pattern consistent with sink-source relationships. These experiments were repeated in triplicate with similar results.

### SMART leaf-specific phenotyping

During the course of these experiments, we noticed that within treatments, the color of individual leaves within the *Arabidopsis* rosette changed substantially, in a manner consistent with source-sink relationships (Figure 4C). While these Fe-associated leaf-specific phenotypes have been described at the molecular level (Schuler *et al.*, 2012), segmentation and quantification of visible traits in a leaf-specific manner is not standard in current phenotyping pipelines. Hence, we implemented a watershed segmentation algorithm to segment individual leaves within the *Arabidopsis* rosette (Figure 5). This watershed segmentation algorithm first uses *Euclidean Distance Transform* to calculate the Euclidean distance map to the closest zero (*i.e*., background pixel) for each of the foreground pixels. In the following step, each peak in the Euclidean distance map was used as a marker for the watershed segmentation to identify and count each of the leaves within the rosette (Figure 5B). Once individual leaves were isolated, all different traits described for the full rosette were extracted including color using the dominant clustering method described before (Table 1; Supplementary Table 2, and Figure 5C-D).

**Figure 5.**
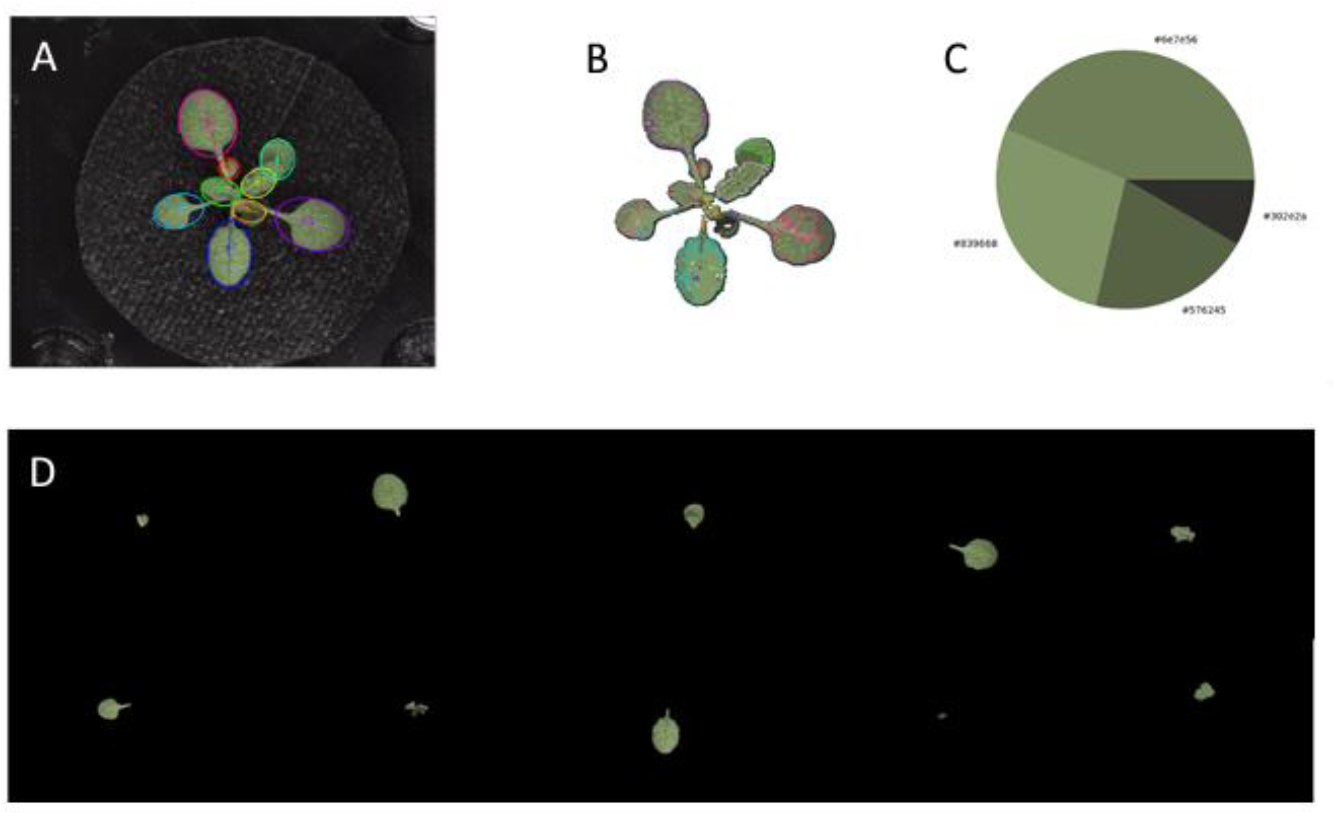
Leaf-specific segmentation of Arabidopsis rosette using SMART. (a-b) SMART is capable of segmenting individual leaves through watershed segmentation. (c-d) color is described as the top four dominant colors using hexadecimal values, and once the individual leaves have been identified and extracted, all the morphological traits described in Table 1 can be quantified in a leaf-specific manner.

To test the capabilities of our leaf-specific phenotyping approach, we compared plants grown in standard Zn levels (1 μM) plus Fe (1 μM Zn+Fe) with plants grown in 10 μM Zn without Fe (10 μM Zn-Fe). We chose these two conditions as starting point based on the data obtained using whole rosette phenotyping (Figure 4), where color changes in a leaf-specific manner were consistently observed. In turn, our individual leaf segmentation approach allowed us to classify individual rosette leaves in three main clusters (Figure 6). Cluster 1 represents the first true leaves within the rosette, whose area remain constant through the progression of the experiment; however, and as excepted for a *source* leaf, these leaves receive and retain the majority of micronutrients coming through the xylem stream and dictate the distribution of micronutrient across the rosette in a pattern consistent with their orthostichy (Khan *et al.*, 2018). Despite minimal changes in individual leaf area, Cluster 1 leaves show clear differences in color, and under the experimental conditions imposed, leaves from plants grown with 10 μM Zn-Fe display hexadecimal color values consistent with darker green compared to the same leaves from plants grown in 1 μM Zn+Fe (Figure 6B and C). In contrast, Cluster 2 leaves represent individual leaves transitioning from sink (developing leaves) to source leaves (fully photosynthetic). In this cluster, the leaf area and color became more dependent on the environmental conditions (*i.e*., nutrient availability) and their unique and autonomous developmental program. For instance, in 1 μM Zn+Fe, the leaf area of Cluster 2 leaves increased over time and their color remain constant through the entire experiment. However, growth of Cluster 2 leaves from plants grown in 10 μM Zn-Fe was variable, with some leaves being larger than plants grown with 1 μM Zn+Fe, while others remain smaller (Figure 6B). Moreover, capturing color throughout the experiment allowed us to observe and quantify the transition of Cluster 2 leaves from dark green to yellow and this transition was consistent with the availability of high Zn and Fe at the beginning of the experiment to lack of Fe in the later days of the experiment. (Figure 6C). Lastly, Cluster 3 leaves represent mostly *sink* leaves; thus, their appearance, color and size were more dependent on the nutrient availability. For instance, Cluster 3 leaves appearance was delayed in plants grown in 10 μM Zn-Fe compared to plants grown in 1 μM Zn+Fe (Figure 6A-B). Moreover, Cluster 3 leaves from plants grown in 10 μM Zn+Fe were more chlorotic compared to leaves from plants grown in 1 μM Zn+Fe (Figure 6C). Altogether, these results demonstrate that *OPEN leaf* and SMART were able to capture the dynamic behavior of individual leaves within the *Arabidopsis* rosette when plants were exposed to different levels of nutrient availability.

**Figure 6.**
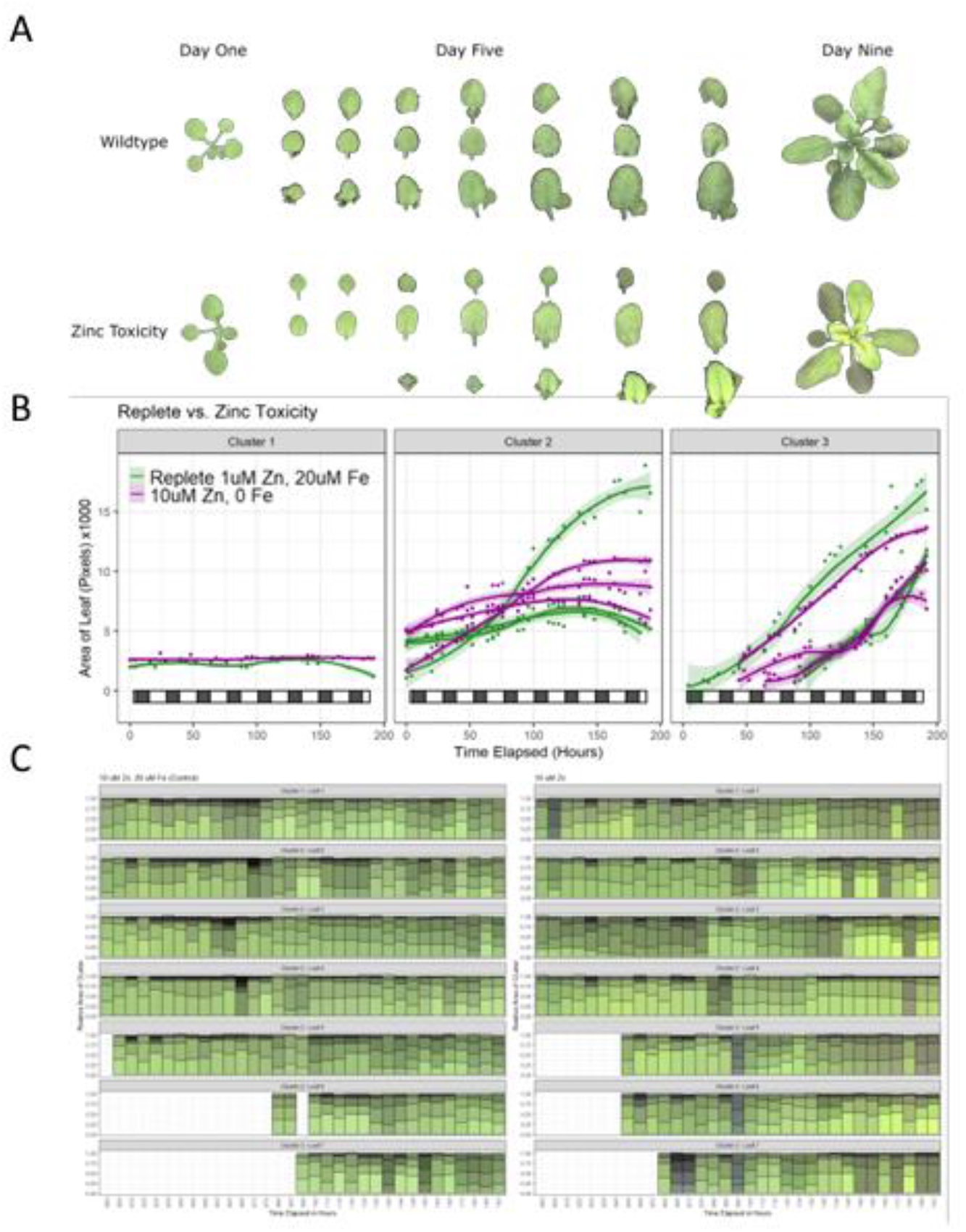
Nutrient availability dictates the fate of individual leaves within the Arabidopsis rosette. (a) the *OPEN leaf* – SMART pipeline is able to track individual leaves during plant growth and development. (b) Changes in the Fe/Zn ratio have a severe impact on individual leaf size and color and leaf clustering was consistent with the leaf developmental stage (source or sink). (c) Individual color clustering of the 4 dominant colors allowed the dynamic tracking of color transitioning during the entire experiment a at leaf-specific resolution. Each bar plot represents the proportional distribution of each dominant color in individual leaves. The plots are representative of one experiment and similar results were obtained in three independent experiments.

## Discussion

The Arabidopsis genome was released in the year 2000 and after more than two decades of intensive molecular research, 30% of its 25,000+ genes remain without an assigned function and a greater number have been annotated based on homology without experimental evidence of their putative function (Wang *et al.*, 2019). There are many factors behind this gap in knowledge, including gene lethality or redundancy within gene families, which is not uncommon in higher plants. In addition, many mutations lead to subtle or transient phenotypes which are difficult to reproduce and capture with standard end-point documentation methods (*i.e*., photos at the end of an experiment) or subject to experimental bias from environmental conditions or image processing protocols such as manual extraction of leaf traits including shape, size and color (Dobrescu *et al.*, 2017; Zhang *et al.*, 2020; Hernández-Herrera *et al.*, 2021).

High throughput plant phenotyping (HTPP) has emerged as a technology to solve many of these issues including consistency during data acquisition and processing (Acosta-Gamboa *et al.*, 2016; Vasseur *et al.*, 2018) HTPP has facilitated the discovery of novel genes and gene function through the use of large mutant collections and diversity panels or ecotypes (Campos *et al.*, 2021; Xie *et al.*, 2021). Unfortunately, commercial HTPP devices remain prohibitively expensive and thus are limited to the very few labs that can afford them. Many open-source HTPP devices have also been described but their assembly and operation require significant expertise in mechanical engineering and computer science, which is not standard nor common in plant biology departments. However, a full and comprehensive characterization of the *dark genome* will require a collective effort within the plant community and at the core of this efforts is accessibility to HTPP technologies and the use of standard practices to conduct experiments under well-defined environmental conditions. This *Internet of Phenotyping* (IoP) in which creating an integrated network of HTPP technologies, user-interactions, and image processing can greatly accelerate development in HTPP.

Here we described *OPEN leaf*, an open-source HTPP device with cloud connectivity and remote bilateral communication with users (Figures 1 and 2). *OPEN leaf* design is user-centered and takes advantage of affordable materials available worldwide, with an approximate cost of $1000 USD (as of Dec 2021; Manual assembly available in supplementary data). *OPEN leaf* together with the SMART processing suite offers a one-stop solution for HTPP of *Arabidopsis* rosettes, from continuous and automatic data collection to data storage, sharing and processing. The cloud integration solves one of the main issues in HTP approaches known as the FAIR principle for data management, where HTP data should be ideally findable, available, identifiable, and reusable for reasonable scientific advancement (Wilkinson *et al.*, 2016). The modular structure of *OPEN leaf* allows users to capture images at a user-defined frequency without the need for cloud or internet connectivity. This modularity also allows users to incorporate additional sensors such as NIR, fluorescence, multi- and hyperspectral cameras; however, each of these sensors will proportionally increase the cost of the HTPP device and data output.

In this work, we also aimed to demonstrate that despite being one of the most basic sensors, visible cameras and RGB images, are still underutilized and several novel phenotypes occurring within the rosette of *Arabidopsis* are yet to be identified, quantified, and characterized. For instance, rosette growth and development is known to be controlled by the intrinsic genetic program of the plant but can also be modified by environmental factors such as nutrient availability or the presence of pathogens (Hindt & Guerinot, 2012; Kim *et al.*, 2014). In turn, our whole-rosette phenotyping pipeline was able to document the growth delay of plants experiencing Fe deficiency (Figure 3). Growth delay in Arabidopsis roots has been found to be mediated by the gibberellin pathway and the corresponding DELLA repressor proteins (Wild *et al.*, 2016). For instance, Arabidopsis plants lacking all DELLA proteins have longer roots during Fe deficiency compared to wildtype plants, suggesting that growth arrest during Fe deficiency is the result of proper Fe sensing followed by downstream hormone-developmental events but not the result of energy shortages due to impaired photosynthesis and respiration. Notably, Fe sensing at the whole plant level is sensed in source leaves, which communicate the Fe status of the plant to roots *via* the phloem to either increase or repressed Fe uptake at the root level (Khan *et al.*, 2018). It is therefore expected that a similar hormone-developmental signaling pathway associated with Fe sensing in leaves takes place; however, the molecular mechanisms behind Fe sensing in leaves remain largely unknown. In turn, our phenotyping system was capable of providing detailed dynamic information about the timing of Fe-deficiency responses in leaves thus offering researchers the opportunity to select ideal time windows for further in-depth molecular analyses (Figure 3).

Hydroponic experiments have provided a unique opportunity to explore plant responses to nutrient availability at a very high resolution and our *OPEN leaf*-SMART pipeline captured subtle differences in rosette size and color during experiments were Fe and Zn availabilities were modified (Figure 4). Moreover, our phenotyping pipeline demonstrates that significant changes occur within the *Arabidopsis* rosette, in a leaf-specific manner, when the ratio of Fe/Zn availability changes (Figure 5). Leaf responses to nutrient availability are expected to be striking different from roots as the *Arabidopsis* rosette includes both sink and source tissues (*i.e*., young and mature leaves), while roots are by definition a sink tissue. In fact, unique leaf-specific transcriptional responses have been described during Fe deficiency experiments in *Arabidopsis* where prolonged Fe deprivation leads to a restricted distribution of Fe to developing (sink) leaves to preserve optimal photosynthesis in mature (source) leaves (Hantzis *et al.*, 2018; McInturf *et al.*, 2021).

While the precise molecular mechanisms behind Fe economy reprograming is ongoing, our *OPEN leaf*-SMART phenotyping pipeline was able to capture the timing of such leaf-specific changes, thus offering the sought-after time window to pursue additional molecular analyses to unravel the mechanisms behind plant responses and adaptation to nutrient availability in a leaf-specific manner.

## Supporting information

STL files for Assembly

## Acknowledgments

This research was supported in part by the US National Science Foundation (MCB-1818312 and IOS-1734145 to DMC), the MU Honors College and Peggy and Andrew Cherng Charitable Foundation (LS), the USDOE ARPA-E ROOTS (DE-AR0000821), the NSF CAREER Award No. 1845760 to A.B, and the US National Institute of Health BD2K Training Grant T32HG009060 grant (STK). We would also like to thank Wesley Paul Bonelli for his contribution to design the script collection data Any opinions findings, and conclusions or recommendations expressed in this material are those of the author(s) and do not necessarily reflect those of ARPA-E or the National Science Foundation.

## Author Contribution

LGS, SM and DM conceived the rpoject, LGD and SM designed HTPP device. EW performed plant experiments on the system. SL and AB developed the SMART imaging pipeline. LGS, SK and DM worked on data interpretation and visualization. LS and DM wrote the manuscript.

## Data Availability

The data that supports the findings of this study are available in the supplementary material of this article

## Supplementary data

Manual assembly

STL files (floaters and camera assembly)

**Supplementary Table 1** Rosette traits extracted using SMART.

**Supplementary Table 2** Leaf-specific traits extracted using SMART.

## References

Acosta-Gamboa, L, Liu S, Langley E, Campbell Z, Guerrero N, Mendoza Cozatl D, Lorence A. 2016. Moderate to severe water limitation differentially affects the phenome and ionome of Arabidopsis. Functional Plant Biology 44.

Awlia, M, Nigro A, Fajkus J, Schmoeckel SM, Negrão S, Santelia D, Trtílek M, Tester M, Julkowska MM, Panzarová K. 2016. High-Throughput Non-destructive Phenotyping of Traits that Contribute to Salinity Tolerance in Arabidopsis thaliana. Frontiers in Plant Science 7: 1414.

Billiau, K, Sprenger H, Schudoma C, Walther D, Köhl KI. 2012. Data management pipeline for plant phenotyping in a multisite project. Functional Plant Biology 39: 948–957.

Buckner, E, Madison I, Chou H, Matthiadis A, Melvin CE, Sozzani R, Williams C, Long TA. 2019. Automated Imaging, Tracking, and Analytics Pipeline for Differentiating Environmental Effects on Root Meristematic Cell Division. Frontiers in Plant Science 10: 1487.

Campos, ACAL, van Dijk WFA, Ramakrishna P, Giles T, Korte P, Douglas A, Smith P, Salt DE. 2021. 1,135 ionomes reveal the global pattern of leaf and seed mineral nutrient and trace element diversity in Arabidopsis thaliana. The Plant Journal 106: 536–554.

Des Marais DL, Hernandez KM, Juenger TE. 2013. Genotype-by-Environment Interaction and Plasticity: Exploring Genomic Responses of Plants to the Abiotic Environment. Annual Review of Ecology, Evolution, and Systematics 44: 5–29.

Dobrescu, A, Scorza LCT, Tsaftaris SA, McCormick AJ. 2017. A “Do-It-Yourself” phenotyping system: measuring growth and morphology throughout the diel cycle in rosette shaped plants. Plant Methods 13: 95.

Flood, PJ, Kruijer W, Schnabel SK, van der Schoor R, Jalink H, Snel JFH, Harbinson J, Aarts MGM. 2016. Phenomics for photosynthesis, growth and reflectance in Arabidopsis thaliana reveals circadian and long-term fluctuations in heritability. Plant methods 12: 14–14.

Hantzis, LJ, Kroh GE, Jahn CE, Cantrell M, Peers G, Pilon M, Ravet K. 2018. A Program for Iron Economy during Deficiency Targets Specific Fe Proteins. Plant Physiology 176: 596–610.

van der Heijden G, Song Y, Horgan G, Polder G, Dieleman A, Bink M, Palloix A, van Eeuwijk F, Glasbey C. 2012. SPICY: towards automated phenotyping of large pepper plants in the greenhouse. Functional Plant Biology 39: 870–877.

Hernández-Herrera P, Ugartechea-Chirino Y, Torres-Martínez HH, Arzola AV, Chairez-Veloz JE, García-Ponce B, Sánchez M de la P, Garay-Arroyo A, Álvarez-Buylla ER, Dubrovsky JG, et al. 2021. Live Plant Cell Tracking: Fiji plugin to analyze cell proliferation dynamics and understand morphogenesis. Plant Physiology: kiab530.

Hindt, MN, Guerinot ML. 2012. Getting a sense for signals: regulation of the plant iron deficiency response. Biochimica et biophysica acta 1823: 1521–1530.

Jackson, SA, Iwata A, Lee S-H, Schmutz J, Shoemaker R. 2011. Sequencing crop genomes: approaches and applications. The New phytologist 191: 915–925.

Khan, MA, Castro-Guerrero NA, McInturf SA, Nguyen NT, Dame AN, Wang J, Bindbeutel RK, Joshi T, Jurisson SS, Nusinow DA, et al. 2018. Changes in iron availability in Arabidopsis are rapidly sensed in the leaf vasculature and impaired sensing leads to opposite transcriptional programs in leaves and roots. Plant, Cell & Environment 41: 2263–2276.

Kim, SH, Son GH, Bhattacharjee S, Kim HJ, Nam JC, Nguyen PDT, Hong JC, Gassmann W. 2014. The Arabidopsis immune adaptor SRFR1 interacts with TCP transcription factors that redundantly contribute to effector-triggered immunity. The Plant Journal 78: 978–989.

Le Marié C, Kirchgessner N, Marschall D, Walter A, Hund A. 2014. Rhizoslides: paper-based growth system for non-destructive, high throughput phenotyping of root development by means of image analysis. Plant Methods 10: 13.

Legris, M, Klose C, Burgie ES, Rojas CCR, Neme M, Hiltbrunner A, Wigge PA, Schäfer E, Vierstra RD, Casal JJ. 2016. Phytochrome B integrates light and temperature signals in *Arabidopsis*. Science 354: 897.

Lopatkin, AJ, Collins JJ. 2020. Predictive biology: modelling, understanding and harnessing microbial complexity. Nature Reviews Microbiology 18: 507–520.

Lundberg, DS, Lebeis SL, Paredes SH, Yourstone S, Gehring J, Malfatti S, Tremblay J, Engelbrektson A, Kunin V, Del Rio TG, et al. 2012. Defining the core Arabidopsis thaliana root microbiome. Nature 488: 86–90.

Maes, WH, Steppe K. 2019. Perspectives for Remote Sensing with Unmanned Aerial Vehicles in Precision Agriculture. Trends in Plant Science 24: 152–164.

Mathieu, L, Lobet G, Tocquin P, Périlleux C. 2015. “Rhizoponics”: a novel hydroponic rhizotron for root system analyses on mature Arabidopsis thaliana plants. Plant Methods 11: 3.

McInturf, SA, Khan MA, Gokul A, Castro-Guerrero NA, Höhner R, Li J, Marjault H-B, Fichman Y, Kunz H-H, Goggin FL, et al. 2021. Cadmium interference with iron sensing reveals transcriptional programs sensitive and insensitive to reactive oxygen species. Journal of Experimental Botany: erab393.

Moore, C, Johnson L, Kwak I-Y, Livny M, Broman K, Spalding E. 2013. High-Throughput Computer Vision Introduces the Time Axis to a Quantitative Trait Map of a Plant Growth Response. Genetics 195.

Nguyen, NT, McInturf SA, Mendoza-Cózatl DG. 2016. Hydroponics: A Versatile System to Study Nutrient Allocation and Plant Responses to Nutrient Availability and Exposure to Toxic Elements. Journal of visualized experiments : JoVE: 54317.

Reynolds, D, Baret F, Welcker C, Bostrom A, Ball J, Cellini F, Lorence A, Chawade A, Khafif M, Noshita K, et al. 2019. What is cost-efficient phenotyping? Optimizing costs for different scenarios. The 4th International Plant Phenotyping Symposium 282: 14–22.

Sadeghi-Tehran, P, Sabermanesh K, Virlet N, Hawkesford MJ. 2017. Automated Method to Determine Two Critical Growth Stages of Wheat: Heading and Flowering. Frontiers in Plant Science 8: 252.

Schuler, M, Rellán-Álvarez R, Fink-Straube C, Abadía J, Bauer P. 2012. Nicotianamine functions in the Phloem-based transport of iron to sink organs, in pollen development and pollen tube growth in Arabidopsis. The Plant cell 24: 2380–2400.

Shendure, J, Ji H. 2008. Next-generation DNA sequencing. Nature Biotechnology 26: 1135–1145.

Tsaftaris, SA, Scharr H. 2019. Sharing the Right Data Right: A Symbiosis with Machine Learning. Trends in Plant Science 24: 99–102.

Vasseur, F, Bresson J, Wang G, Schwab R, Weigel D. 2018. Image-based methods for phenotyping growth dynamics and fitness components in Arabidopsis thaliana. Plant Methods 14: 63.

van der Walt S, Schönberger JL, Nunez-Iglesias J, Boulogne F, Warner JD, Yager N, Gouillart E, Yu T, scikit-image contributors. 2014. scikit-image: image processing in Python. PeerJ 2: e453–e453.

Wang, W, Zhang X, Niittylä T. 2019. OPENER Is a Nuclear Envelope and Mitochondria Localized Protein Required for Cell Cycle Progression in Arabidopsis. The Plant cell 31: 1446–1465.

White, JW, Andrade-Sanchez P, Gore MA, Bronson KF, Coffelt TA, Conley MM, Feldmann KA, French AN, Heun JT, Hunsaker DJ, et al. 2012. Field-based phenomics for plant genetics research. Field Crops Research 133: 101–112.

Wild, M, Davière J-M, Regnault T, Sakvarelidze-Achard L, Carrera E, Lopez Diaz I, Cayrel A, Dubeaux G, Vert G, Achard P. 2016. Tissue-Specific Regulation of Gibberellin Signaling Fine-Tunes Arabidopsis Iron-Deficiency Responses. Developmental Cell 37: 190–200.

Wilkinson, MD, Dumontier M, Aalbersberg IjJ, Appleton G, Axton M, Baak A, Blomberg N, Boiten J-W, da Silva Santos LB, Bourne PE, et al. 2016. The FAIR Guiding Principles for scientific data management and stewardship. Scientific Data 3: 160018.

Xiao, Y, Liu H, Wu L, Warburton M, Yan J. 2017. Genome-wide Association Studies in Maize: Praise and Stargaze. Molecular plant 10: 359–374.

Xie, J, Fernandes SB, Mayfield-Jones D, Erice G, Choi M, E Lipka A, Leakey ADB. 2021. Optical topometry and machine learning to rapidly phenotype stomatal patterning traits for maize QTL mapping. Plant Physiology 187: 1462–1480.

Xu Y. 2016. Envirotyping for deciphering environmental impacts on crop plants. Theoretical and Applied Genetics 129: 653–673.

Yang, W, Feng H, Zhang X, Zhang J, Doonan JH, Batchelor WD, Xiong L, Yan J. 2020. Crop Phenomics and High-Throughput Phenotyping: Past Decades, Current Challenges, and Future Perspectives. Molecular Plant 13: 187–214.

Yang, W, Guo Z, Huang C, Duan L, Chen G, Jiang N, Fang W, Feng H, Xie W, Lian X, et al. 2014. Combining high-throughput phenotyping and genome-wide association studies to reveal natural genetic variation in rice. Nature Communications 5: 5087.

Zhang, X, Hu Z, Guo Y, Shan X, Li X, Lin J. 2020. High-efficiency procedure to characterize, segment, and quantify complex multicellularity in raw micrographs in plants. Plant Methods 16: 100.

